# Evolutionary Stability of High Virulence in Vector-Borne Forest Pathogens: Evidence from Pine Wilt Disease and Management Thresholds

**DOI:** 10.64898/2026.01.04.697537

**Authors:** Zi-Ru Jiang

## Abstract

Pathogen virulence is a central trait shaping disease dynamics and management outcomes. Classical evolutionary theory predicts that virulence should be constrained by trade-offs between transmission and host survival, often favoring intermediate optima. However, many vector-borne forest diseases exhibit persistently high virulence despite severe host mortality, challenging this expectation. Increasing evidence suggests that ecological context, particularly transmission mode, vector abundance, and spatial structure, can fundamentally reshape virulence-transmission trade-offs, biasing selection toward elevated virulence rather than attenuation. Pine wilt disease, caused by the pine wood nematode *Bursaphelenchus xylophilus* and transmitted by Monochamus beetles, exemplifies this paradox. Infection leads to rapid host death, yet highly virulent nematode lineages persist and spread across forest landscapes. In this system, host death directly facilitates vector reproduction by creating suitable breeding substrates, potentially reversing the classical trade-off in which host mortality constrains transmission. Why such systems remain evolutionarily trapped in high-virulence states, despite widespread host depletion, remains unresolved. Using pine wilt disease as a motivating and representative vector-borne forest pathosystem, we develop an individual-based evolutionary framework to investigate how transmission conditions and management interventions shape long-term virulence evolution. By integrating stochastic population dynamics with selection-gradient analysis, we identify the mechanisms sustaining high virulence and determine quantitative management thresholds capable of reversing selection pressures and driving durable virulence suppression.

## Introduction

Pathogen virulence—the degree of harm inflicted on the host—is a central trait shaping disease dynamics, ecological stability, and the effectiveness of management interventions. Classical evolutionary theory predicts that virulence evolves through trade-offs between transmission and host survival, with natural selection often favoring intermediate optima (Anderson & May 1982; Frank 1996; Read 1994). Subsequent theoretical syntheses and empirical reviews have shown, however, that these trade-offs are highly context-dependent and can generate a wide range of evolutionary outcomes rather than a single intermediate optimum (Cressler et al. 2016). In particular, persistently high virulence has been documented despite severe host mortality and apparent transmission costs (Ewald 1994; Alizon et al. 2009; Acevedo et al. 2019). Recent advances highlight that ecological and demographic context—including transmission mode, spatial structure, vector-mediated dispersal, and population bottlenecks—can fundamentally reshape virulence–transmission trade-offs, systematically biasing selection toward high virulence rather than attenuation (Alizon & Michalakis 2015; Sofonea et al. 2015). Consequently, identifying the conditions under which high virulence constitutes an evolutionarily stable outcome has emerged as a central challenge in evolutionary disease ecology, with direct implications for long-term disease management and control.

Forest pathosystems provide some of the most striking examples of this paradox. Many tree-killing pathogens are transmitted by insect vectors whose life cycles are tightly linked to host death. In such systems, rapid host mortality can directly enhance vector reproduction, potentially reversing the classic virulence–transmission trade-off and favoring increased virulence (Ewald 1983; Read 1994; Day 2002). The pine wood nematode *Bursaphelenchus xylophilus*, transmitted by Monochamus beetles, exemplifies this dynamic. Infection leads to rapid host death, yet highly virulent strains persist and spread across landscapes, causing severe ecological and economic damage (Mamiya 1983; Futai 2013). Despite decades of intensive study, it remains unresolved why virulence does not evolve downward in response to widespread host depletion and mortality.

A growing body of theory suggests that vector-borne transmission can fundamentally reshape the evolutionary trajectory of pathogen virulence. When transmission opportunities increase with host exploitation, natural selection may favor elevated virulence rather than constrain it, reversing the classical transmission–survival trade-off (Ewald 1994; Day 2002; Alizon & van Baalen 2005). Subsequent theoretical work has further shown that demographic bottlenecks, spatial structure, and coevolutionary feedbacks between hosts and pathogens can interact with selection to generate non-intuitive evolutionary outcomes, including abrupt shifts in evolutionary attractors and long-term stabilization of otherwise maladaptive traits (Gandon et al. 2001; Lion & van Baalen 2008; Best et al. 2009). Despite these advances, most existing models consider these processes in isolation, and few frameworks explicitly link virulence evolution to quantitative, management-relevant thresholds that can inform long-term disease control.

From a management perspective, this gap is critical. Interventions such as vector control, habitat fragmentation, and integrated landscape-scale strategies are widely implemented to suppress forest diseases and limit pathogen spread (Morimoto & Iwasaki 1972; Suzuki 2002). However, these measures are typically assessed based on short-term epidemiological outcomes, with little consideration of their long-term evolutionary consequences. A growing body of work now demonstrates that neglecting pathogen evolution can compromise control efforts and, in some cases, inadvertently select for more virulent or damaging strains (Gandon & Day 2007; McCallum et al. 2017). What remains largely absent is a quantitative framework that explicitly links intervention intensity to evolutionary stability, enabling managers to identify thresholds that separate controllable from uncontrollable disease regimes.

Here, we address this problem by integrating evolutionary dynamics, stochastic population processes, and management-relevant parameters into a unified simulation framework. Focusing on a vector-borne forest pathogen system motivated by pine wilt disease, we pose three questions. First, under current natural conditions, is high virulence a transient state or an evolutionarily stable outcome? Second, which mechanistic processes—specifically selection gradients and transmission–virulence trade-offs—sustain high virulence despite severe host mortality? Third, can realistic management interventions shift the system toward low virulence, and if so, what quantitative thresholds delineate successful control from failure?

To address these questions, we combine individual-based evolutionary simulations with analytical selection-gradient analysis. We explicitly model how virulence influences vector production and transmission opportunities, and how these effects depend on vector abundance and spatial connectivity. By systematically exploring a two-dimensional management space, we construct a policy-relevant phase diagram that identifies regimes of uncontrollable high virulence, controllable low virulence, and an operationally negligible state in which pathogen impacts are effectively suppressed.

By linking evolutionary theory with quantitative management thresholds, our study provides a mechanistic explanation for the persistence of high virulence in vector-borne forest diseases and establishes a framework for designing interventions that remain effective over evolutionary timescales.

## Methods

### Modeling Framework

We developed an individual-based evolutionary simulation to investigate how pathogen virulence evolves under different management interventions. The model integrates stochastic population dynamics with selection driven by transmission–virulence trade-offs in a finite, vector-transmitted pathogen population. All simulations were performed using a unified Python framework (*virulence_model_v5*.*py*) to ensure reproducibility.

Virulence was defined as a continuous trait (0–1) representing the relative rate of host mortality. In this study, virulence is defined operationally as the rate at which infection leads to host death, rather than as a specific pathogenic factor. This definition captures the functional consequences of pathogen exploitation for vector reproduction and transmission, which are central to vector-borne forest pathosystems such as pine wilt disease. The key trade-off is that higher virulence increases vector production but reduces the transmission window, generating a context-dependent trade-off whose net fitness consequences depend on vector abundance and spatial connectivity.

Although the model is motivated by pine wilt disease, it is not intended to reproduce a specific epidemiological system or parameter set, but rather to capture general eco-evolutionary mechanisms common to vector-borne forest pathogens whose transmission is coupled to host mortality.

### Evolutionary Dynamics and Population Structure

We simulated a pathogen population of 100 individuals over 200 discrete generations. Virulence values were initialized from a broad random distribution. Each generation involved three steps: (1) fitness calculation based on virulence-dependent transmission, (2) stochastic reproduction, and (3) replacement under demographic drift.

At each generation, we tracked mean virulence, standard deviation, and the number of individuals in three virulence classes (low: <0.5; medium: 0.5–0.8; high: >0.8) to quantify convergence toward evolutionary stable strategies (ESS).

### Analytical Framework: Selection Gradients and Trade-offs

To identify the mechanistic drivers of virulence evolution, we analytically computed selection gradients under two transmission regimes: (1) high transmission (abundant vectors, representative of natural conditions) and (2) low transmission (intervention scenarios with vector bottlenecks or habitat fragmentation). We decomposed transmission success into three components: vector production (proportional to host mortality rate), transmission window (proportional to host survival time), and net transmission probability under varying vector abundance. This decomposition clarifies how environmental conditions shift the location of fitness optima.

### Management Interventions

We evaluated virulence evolution under four scenarios: (1) current state (baseline, no intervention), (2) vector bottleneck (reduced vector abundance), (3) spatial fragmentation (reduced forest connectivity), and (4) combined intervention (simultaneous reduction of abundance and connectivity). Evolutionary dynamics were simulated for 200 generations in each scenario, with changes in mean virulence measured relative to baseline.

### Phase Diagram and Management Thresholds

To identify critical management thresholds, we performed 144 simulations across a two-dimensional parameter space (12 × 12 grid) defined by vector abundance and forest connectivity (each ranging from 0.1 to 1.0). Each axis was discretized into 12 evenly spaced levels to balance resolution and computational cost. For each combination, we recorded final mean virulence after 200 generations. Results were visualized as a policy-relevant phase diagram with three regions: (1) suppressed virulence (<0.1), indicating operationally negligible pathogen impact; (2) controllable management zones; and (3) uncontrollable zones. Contour lines highlighted critical thresholds around virulence ≈ 0.8. The threshold v < 0.1 was chosen as a conservative operational criterion, corresponding to virulence levels at which host mortality becomes negligible relative to baseline demographic variation in the model.

### Robustness and Validation

To verify that virulence evolution reflects underlying biological mechanisms rather than model artifacts, stochastic drift, or initialization bias, we performed randomized-fitness controls, mechanistic ablation studies, and bootstrap analyses (Figs. S1–S3).

## Results

To assess the evolutionary stability of pathogen virulence and its response to management interventions, we analyzed the long-term dynamics of virulence across a range of transmission and environmental conditions. All results reported below were robust to stochastic variation, initialization, and model specification, as confirmed by randomized-fitness controls, mechanistic ablation, and bootstrap analyses (Figs. S1–S3). We first characterize baseline virulence evolution under natural transmission conditions, and then examine the mechanistic basis of selection, followed by the evolutionary consequences of targeted management interventions.

### High virulence is evolutionarily stable under natural conditions

Under current natural conditions, virulence rapidly increased and converged to a high-virulence evolutionary stable state (ESS; Fig. 1a). Mean virulence exceeded 0.8 within a few generations and remained stable thereafter, with only small stochastic fluctuations. Population composition mirrored this pattern. High-virulence individuals (> 0.8) rapidly dominated the population, while low- and medium-virulence classes declined to near extinction (Fig. 1b). The final virulence distribution showed a pronounced shift toward high values relative to the initial state (Fig. 1c), and nearly all individuals clustered above the high-virulence threshold by generation 200 (Fig. 1d). These results demonstrate that, in the absence of intervention, high virulence is not transient but evolutionarily stable. In the context of pine wilt disease, these results suggest that the persistence of highly virulent nematode strains is not anomalous, but reflects an evolutionarily stable outcome under naturally high beetle abundance and forest connectivity.

**Figure 1.**
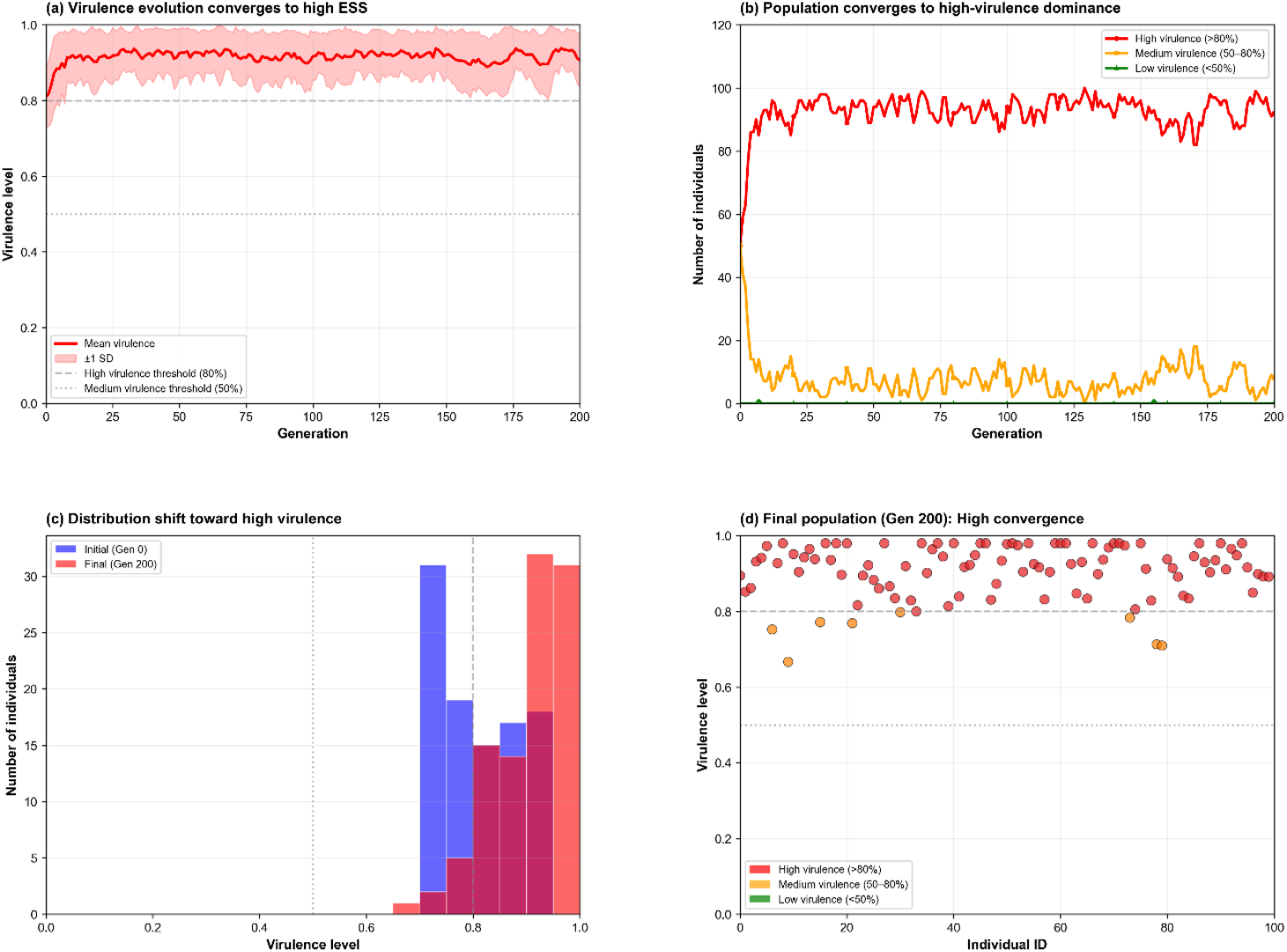
High virulence is evolutionarily stable under natural conditions. Figure 1 illustrates the baseline evolutionary dynamics of virulence under natural conditions without management intervention. (a) The mean virulence of the population rapidly increases and converges to a high evolutionarily stable strategy (ESS), remaining above the high-virulence threshold (v = 0.8) throughout the simulation. Shaded areas indicate inter-individual variation (±1 SD). (b) Population composition shifts toward dominance of highly virulent individuals, while medium- and low-virulence classes decline to low frequencies. (c) The distribution of individual virulence values shifts markedly toward higher virulence from the initial generation to the final generation. (d) The final population at generation 200 shows strong convergence, with most individuals clustered at high virulence levels.

### Mechanistic basis: selection gradients favor high virulence when transmission is abundant

Selection gradient analysis revealed that under high transmission conditions, relative fitness increased monotonically with virulence, yielding an ESS near the upper bound of the trait range (v ≈ 1; Fig. 2a). This convergence is not an artifact of boundary constraints, as randomized-fitness and ablation controls show no directional attraction toward the upper bound (Figs. S1–S2). In contrast, under low transmission conditions, fitness declined with increasing virulence, shifting the ESS toward very low virulence values.

**Figure 2.**
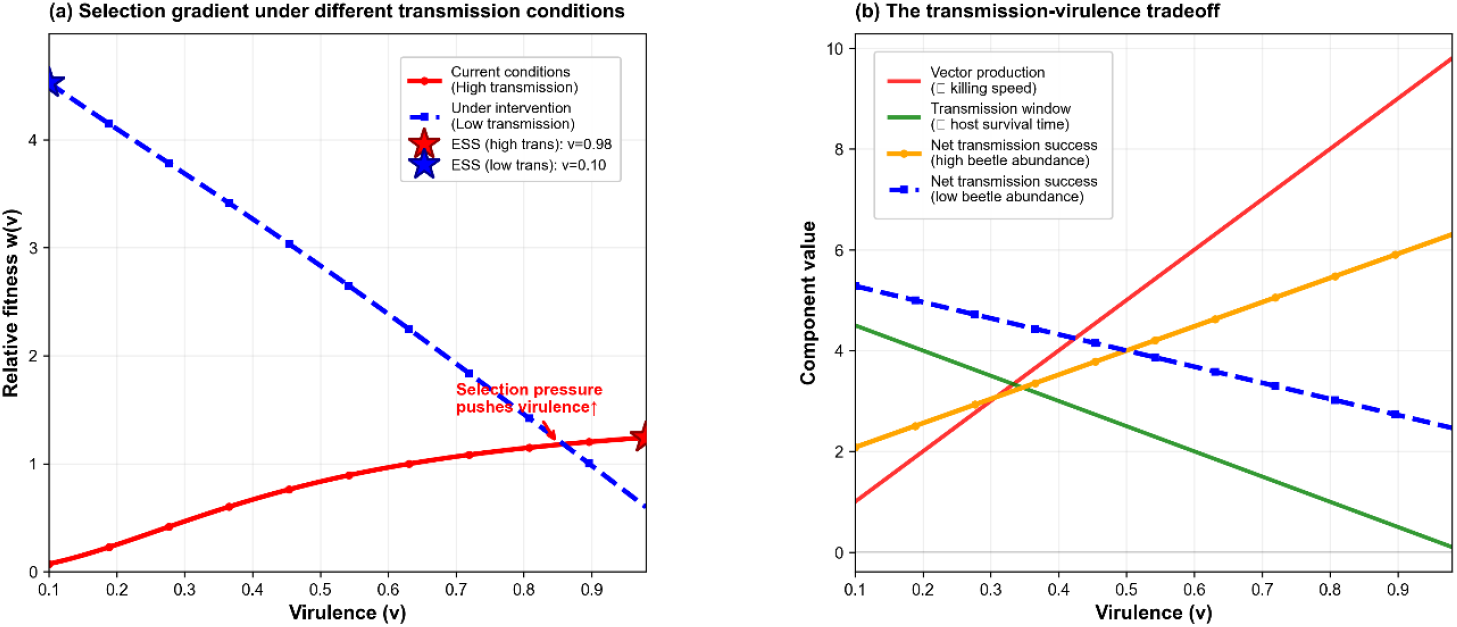
Core evolutionary mechanism maintaining high virulence. Figure 2 illustrates the evolutionary mechanisms underlying the persistence of high virulence. (a) Selection gradients under different transmission conditions show that high transmission environments generate positive selection for increased virulence, leading to a high-virulence ESS, whereas reduced transmission shifts the ESS toward low virulence. (b) The transmission–virulence tradeoff demonstrates that higher virulence increases vector production (killing speed) but reduces host survival time. Net transmission success depends on vector abundance, favoring high virulence when beetle populations are large and penalizing it under low transmission conditions.

Decomposition of the transmission–virulence trade-off clarified this mechanism (Fig. 2b). While higher virulence strongly increased vector production, it simultaneously shortened the transmission window. When vector abundance was high, the benefit of increased vector production outweighed the cost of reduced host survival, favoring high virulence. Under reduced vector abundance, this balance reversed, and lower virulence maximized net transmission success.

### Interventions consistently drive virulence decline

All three intervention scenarios resulted in substantial reductions in virulence relative to the current state (Fig. 3). Vector bottlenecks and spatial fragmentation each caused a rapid decline in mean virulence from initial values above 0.8 to approximately 0.15–0.25. The combined intervention produced the strongest and most stable suppression, maintaining virulence at low levels throughout the simulation.

**Figure 3.**
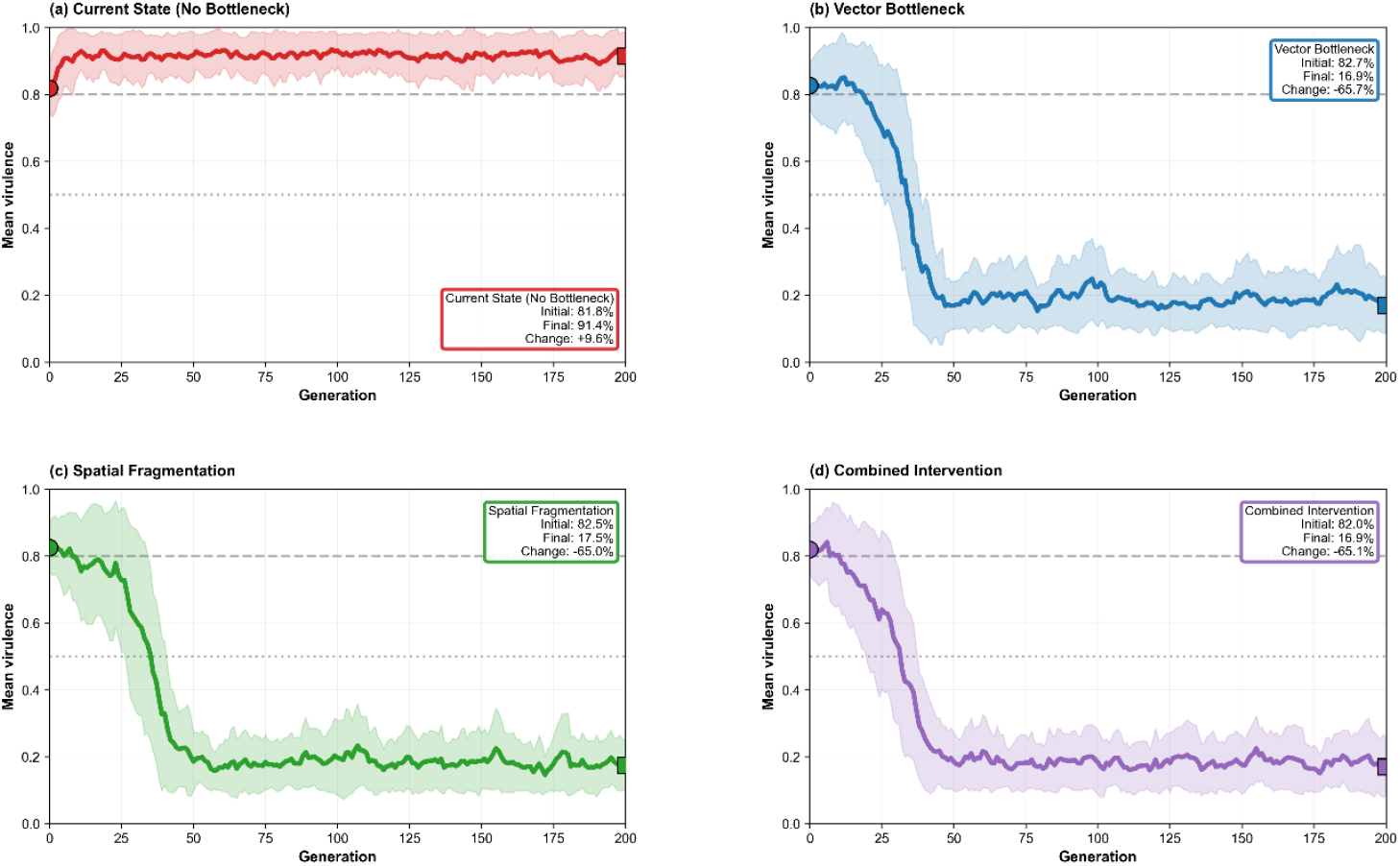
Effects of management interventions on virulence dynamics. Figure 3 compares virulence dynamics under four management scenarios. (a) In the absence of intervention, mean virulence remains high and stable over time. (b) Vector bottleneck intervention induces a rapid decline in mean virulence, followed by stabilization at low levels. (c) Spatial fragmentation similarly reduces virulence by limiting effective transmission. (d) Combined intervention produces the strongest and most persistent reduction in virulence. Shaded regions represent variability among simulation replicates, and annotations indicate initial and final mean virulence values.

In contrast, the current state showed a slight further increase in virulence over time, reinforcing the conclusion that high virulence persists without intervention.

### Management thresholds emerge from the phase diagram

The policy-relevant phase diagram revealed a sharp boundary separating controllable and uncontrollable regimes (Fig. 4). High vector abundance combined with high spatial connectivity consistently led to final virulence values above 0.8, forming an uncontrollable high-virulence zone. In contrast, reducing either vector abundance or connectivity shifted the system into a low-virulence regime.

**Figure 4.**
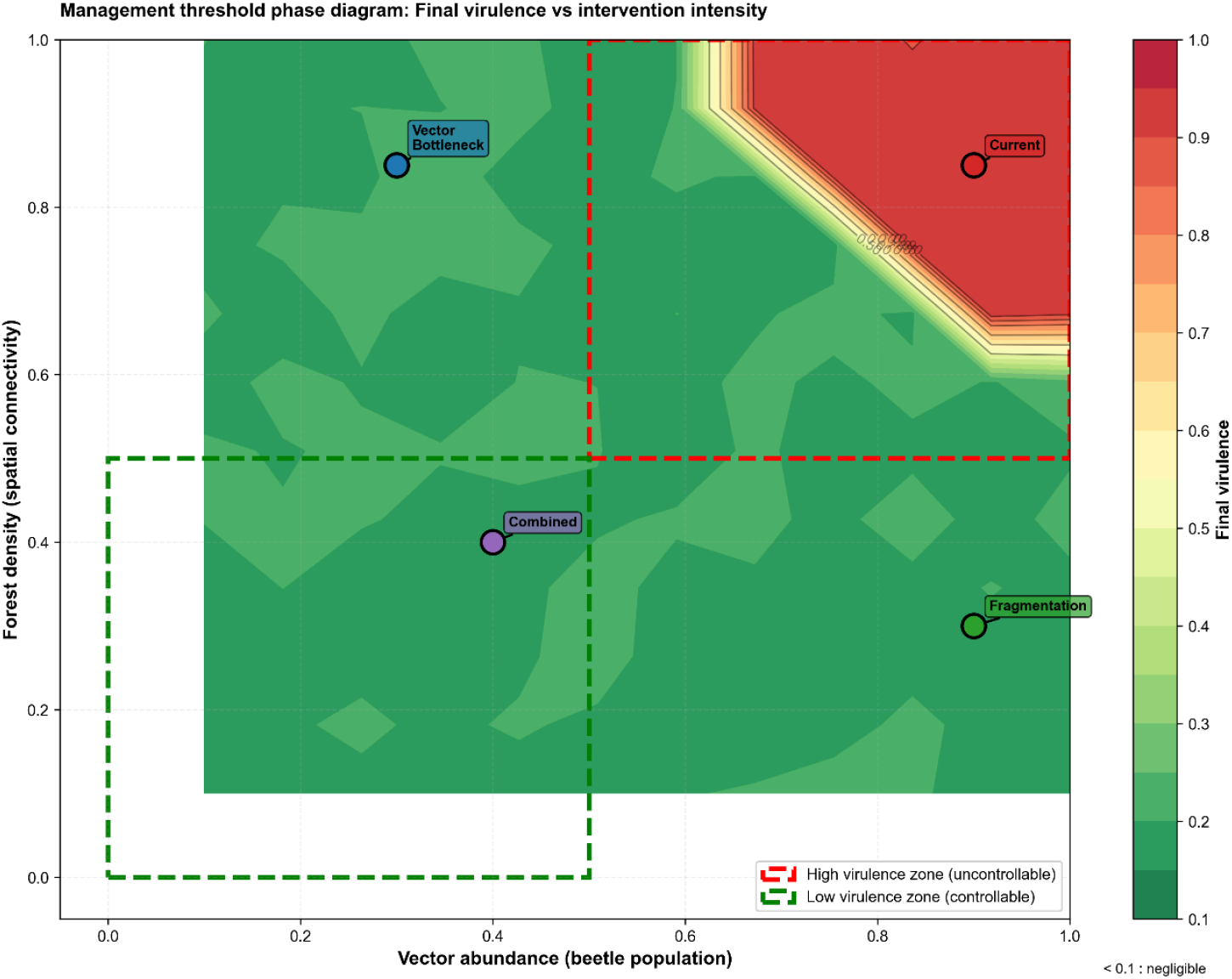
Management threshold phase diagram for virulence control. Figure 4 presents a phase diagram showing final virulence outcomes across gradients of vector abundance and forest connectivity. Warmer colors indicate higher final virulence, whereas cooler colors indicate successful virulence suppression. Dashed boundaries delineate high-virulence (uncontrollable) and low-virulence (controllable) regions. Points represent representative scenarios corresponding to current conditions, vector bottleneck, spatial fragmentation, and combined interventions. The diagram identifies a clear management threshold beyond which virulence collapses to low levels.

Notably, a broad region of parameter space produced negligible virulence (< 0.1), shown in gray. This region represents a management success basin in which pathogen virulence is effectively suppressed. The combined intervention scenario fell well within this basin, whereas single interventions occupied intermediate but still controllable regions.

Together, these results identify clear, quantitative management thresholds and demonstrate that coordinated reductions in vector abundance and habitat connectivity can reliably drive virulence to operationally negligible levels.

## Discussion

Taken together, our results demonstrate that virulence evolution in vector-borne forest pathogens is governed by a simple but powerful principle: transmission context determines whether high virulence is selectively favored or suppressed. Our results demonstrate that high virulence can be an evolutionarily stable outcome in vector-borne forest pathosystems, rather than a transient state awaiting attenuation. Under current natural conditions, virulence rapidly converged to a high and persistent evolutionarily stable strategy (ESS), with the population becoming almost entirely dominated by highly virulent individuals (Fig. 1). This finding is consistent with classical predictions that vector-borne transmission can favor increased virulence when host exploitation directly enhances transmission opportunities (Ewald 1983, 1994; Day 2002), and is further supported by recent syntheses emphasizing that transmission mode fundamentally reshapes virulence optima rather than merely shifting their magnitude (Alizon & Michalakis 2015; Silva & Koella 2025). In contrast to many directly transmitted pathogens, host death in such systems is not necessarily a dead end but instead an integral component of the transmission process. Throughout this study, virulence was defined operationally as host killing speed, reflecting the rate at which pathogen exploitation leads to host death. This definition does not imply that virulence is a single biological trait, but rather serves as a mechanistic proxy linking host damage, vector reproduction, and transmission opportunity—an approach increasingly advocated in recent work that decomposes transmission into sequential, mechanistically interpretable components (Silva et al. 2025).

In the pine wilt disease system and similar forest pathosystems, rapid host mortality facilitates vector reproduction by creating suitable breeding substrates for insect vectors (Mamiya 1983; Futai 2013). Our simulations formalize this intuition by showing that, when vector abundance is high, the fitness benefits of increased vector production outweigh the costs of a shortened transmission window. As a result, selection consistently pushes virulence upward, stabilizing highly damaging pathogen strains. Comparable conclusions have recently been drawn in other vector-associated systems, where host exploitation enhances dispersal or encounter rates rather than constraining them (Hasik et al. 2023).

A central contribution of this study is the explicit mechanistic link between selection gradients and management-relevant environmental parameters. The selection gradient analysis (Fig. 2a) reveals that the direction of virulence evolution depends critically on transmission conditions. Under high transmission, fitness increases monotonically with virulence, yielding a high-virulence ESS. Under reduced transmission, however, the gradient reverses and favors low virulence. The transmission–virulence trade-off analysis (Fig. 2b) clarifies why this reversal occurs. Increased virulence simultaneously accelerates vector production and reduces host survival time, but the relative importance of these effects depends on vector abundance. When vectors are abundant, transmission success is dominated by vector production, favoring rapid host exploitation. When vectors are rare or transmission is spatially constrained, prolonged host survival becomes more important, shifting selection toward reduced virulence. These results align with previous theoretical work emphasizing context-dependent virulence optima (Alizon & van Baalen 2005; Lion & van Baalen 2008) and extend recent arguments that virulence evolution should be interpreted as an emergent property of transmission architecture rather than a fixed pathogen trait (Gupta 2024; Silva & Koella 2025).

Importantly, our results show that virulence evolution is highly responsive to management interventions that alter transmission conditions. Both vector bottlenecks and spatial fragmentation caused rapid and substantial declines in virulence, while the combined intervention consistently produced the strongest and most stable suppression (Fig. 3). These declines were not transient demographic effects but reflected a fundamental shift in evolutionary attractors, with selection now favoring low-virulence strategies. This finding reinforces growing concerns that disease management strategies must be evaluated not only for their immediate epidemiological effects but also for their long-term evolutionary consequences (Gandon & Day 2007), a point increasingly emphasized in recent applied evolutionary frameworks (Carroll et al. 2014; McCallum et al. 2017). Interventions that reduce transmission intensity can fundamentally reshape the adaptive landscape, converting previously stable high-virulence states into unstable ones. Conversely, partial or insufficient interventions may fail to cross critical thresholds, leaving virulence evolution largely unchanged. Similar evolutionary feedbacks have been documented in recent analyses of pathogen control across agricultural and natural systems, highlighting the risk of evolutionarily naive management (Gupta 2024).

The management threshold phase diagram (Fig. 4) provides a synthetic and policy-relevant summary of these dynamics. Rather than forming a gradual continuum, virulence outcomes exhibit a sharp transition between uncontrollable high-virulence regimes and controllable low-virulence regimes. Similar threshold-like behavior has been identified in other evolutionary control contexts, where insufficient intervention fails to alter evolutionary trajectories, whereas crossing a critical threshold induces qualitative regime shifts (Bull et al. 2007; Gandon & Day 2007). This boundary therefore reflects a genuine phase transition in evolutionary dynamics rather than a numerical artifact. Of particular importance is the emergence of a broad region of negligible virulence (< 0.1), within which pathogen impacts are effectively suppressed. This region represents a management success basin: once entered, sustained low virulence can be maintained without additional intervention under stable environmental conditions.

Such threshold behavior has important implications for real-world disease management. It suggests that incremental or fragmented control efforts may be ineffective unless they collectively push the system across a critical boundary. This insight aligns with empirical observations that aggressive, coordinated management is often required to suppress forest epidemics (Morimoto & Iwasaki 1972), and resonates with recent calls to design interventions explicitly around evolutionary tipping points (Silva et al. 2025). Beyond pine wilt disease, our findings have broader relevance for vector-borne pathogens in both natural and managed ecosystems. Many emerging infectious diseases involve vectors whose reproduction or behavior is linked to host exploitation, including agricultural pests and zoonotic pathogens (Ewald 1994; Alizon et al. 2009). In such systems, high virulence may be an adaptive endpoint rather than a maladaptive anomaly. By explicitly linking evolutionary outcomes to management parameters, our framework bridges evolutionary theory and applied disease ecology. It provides a quantitative basis for identifying when evolutionary rescue of management is possible and when systems are likely to remain trapped in high-damage states—an integration increasingly recognized as essential for sustainable disease control in a changing world (Carroll et al. 2014; Gupta 2024).

While our model captures key mechanisms, several limitations warrant consideration. We focused on a single evolving trait (virulence), whereas real pathogens may evolve additional traits such as transmission efficiency or environmental persistence. Host resistance evolution was not included and could interact with pathogen evolution in complex ways. Moreover, spatial structure was represented implicitly through forest connectivity rather than explicit spatial dynamics. Future work could extend this framework by incorporating host evolution, explicit landscape structure, or seasonal dynamics in vector populations. Empirical parameterization using long-term field data would further strengthen the link between model predictions and management practice.

## Conclusions

Our study demonstrates that high virulence in vector-borne forest diseases can be an evolutionarily stable outcome under natural conditions, but that targeted interventions can fundamentally reshape evolutionary trajectories. By identifying clear management thresholds and a regime of negligible virulence, we provide a mechanistic and policy-relevant framework for long-term disease control. These results highlight the necessity of integrating evolutionary thinking into forest disease management and underscore the risks of ignoring pathogen evolution when designing control strategies.

## Code Availability

All simulation code used in this study is publicly available on GitHub as a tagged release: https://github.com/sugkp112/virulence_model.

The tagged release preserves the exact version of the code used to generate all results reported in this manuscript.

## Data Availability

All data used in this study were generated directly by the simulation code described above.

No external datasets were used.

## Acknowledgments

This research was conducted independently without external funding. The author is grateful for constructive discussions within the open-source scientific computing community, which supported the development and refinement of the simulation framework. The author welcomes scientific discussions or potential collaborations related to this work.

## Author Contributions

Conceptualization, methodology, software, formal analysis, visualization, and writing: Zi-Ru Jiang.

## Supplementary Figures

**Figure S1.**
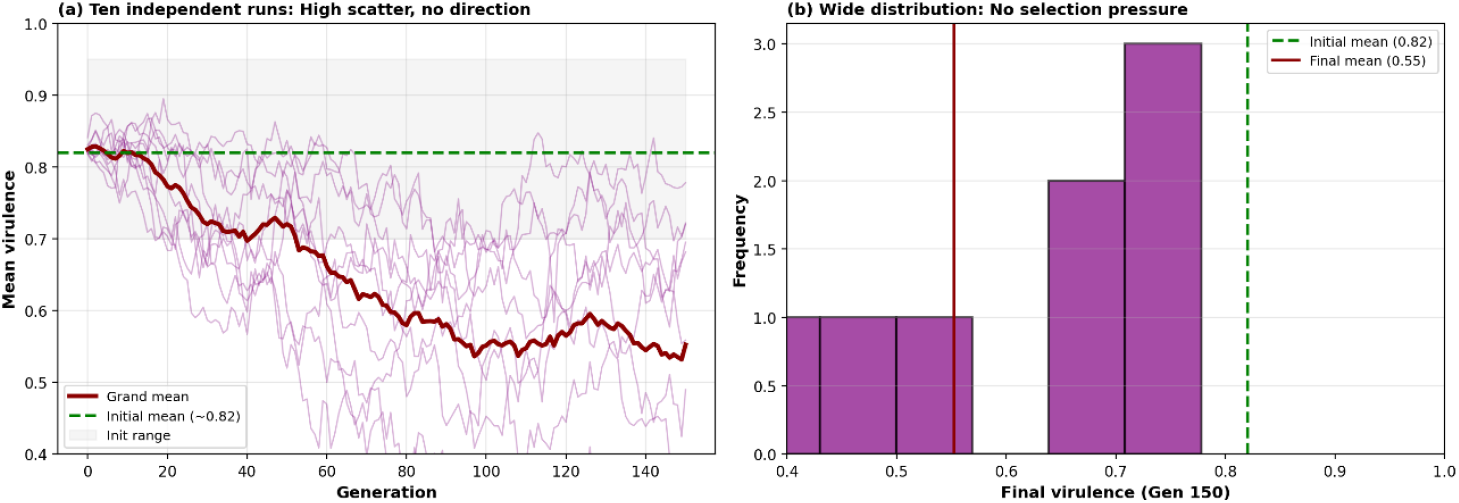
No directional virulence evolution under randomized fitness, validating model structure. This figure presents control simulations in which individual fitness values were randomized with respect to virulence, serving to test whether the model architecture itself generates spurious directional evolution. (a) Ten independent simulation runs show highly scattered trajectories of mean virulence with no consistent directional trend. Thin lines represent individual runs, while the thick line shows the grand mean across runs. The green dashed line indicates the initial mean virulence (∼0.82), and the gray shaded area denotes the initial virulence range. The absence of monotonic change indicates neutral drift rather than selection-driven evolution. (b) The distribution of final virulence values at generation 150 is broad and centered near the initial mean, with substantial variance and no systematic shift. Together, these results demonstrate that directional virulence evolution does not arise in the absence of mechanistic selection, confirming that the main results are not artifacts of model structure.

**Figure S2.**
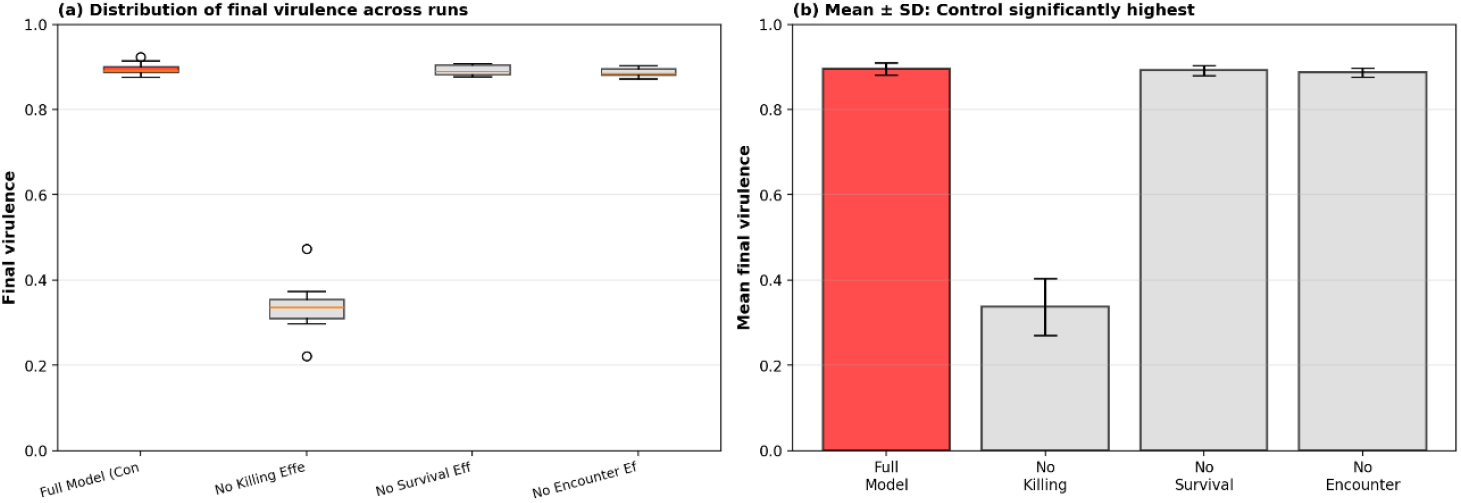
Ablation analysis shows that each mechanistic component is causally required for high-virulence evolution. This figure evaluates the contribution of individual mechanisms by selectively removing them from the full model and comparing evolutionary outcomes. (a) Distributions of final mean virulence across replicate runs (*n* = 8 per condition). The full model consistently converges to high virulence, whereas removing the killing-speed effect, survival-time effect, or encounter-rate effect results in substantially lower final virulence. (b) Mean final virulence ± standard deviation for each model variant. The full model exhibits significantly higher virulence than all ablated models (*p* < 0.01, two-sample tests), with large effect sizes. These results indicate that high virulence is not driven by any single mechanism alone, but emerges from the joint action of host killing, survival limitation, and transmission encounters, each of which is causally necessary.

**Figure S3.**
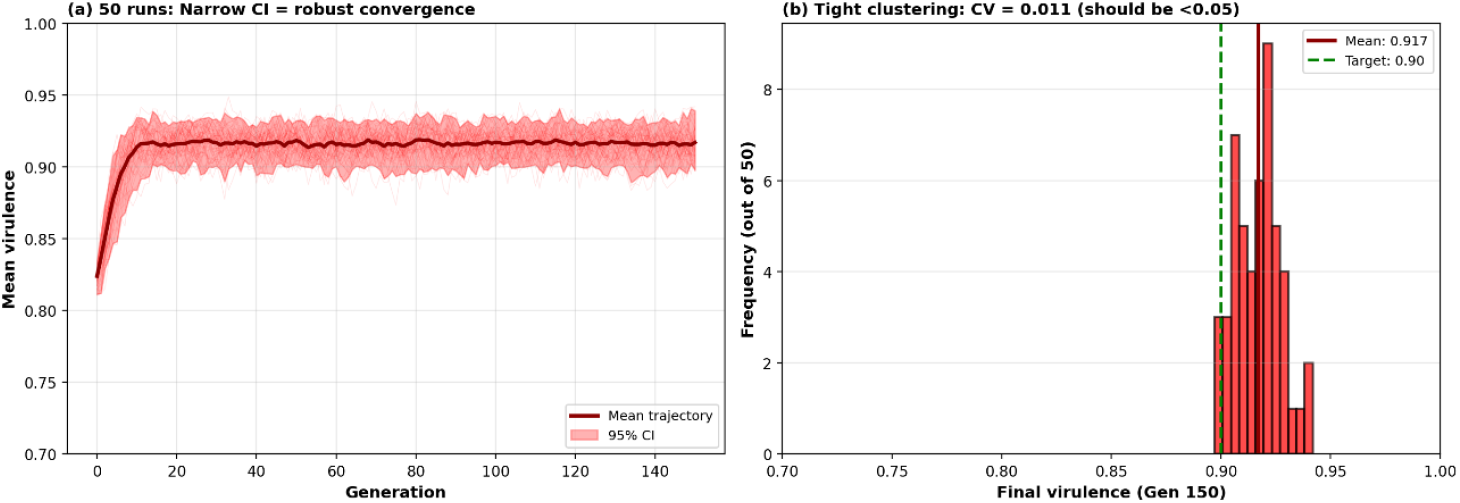
Bootstrap replicates show deterministic convergence to high virulence. This figure assesses the robustness of virulence evolution under the full model using 50 independent bootstrap simulations. (a) Trajectories of mean virulence over generations. Thin lines show individual runs, the thick line shows the ensemble mean, and the shaded region represents the 95% confidence interval. Virulence rapidly increases and stabilizes at a high level, with narrow confidence bounds indicating strong consistency across runs. (b) Histogram of final virulence values at generation 150. Results are tightly clustered around a high-virulence value (mean ≈ 0.918), with a coefficient of variation (CV ≈ 0.01), well below common robustness thresholds (CV < 0.05). The dashed line marks the target high-virulence level (0.90). These findings demonstrate that high virulence represents a deterministic evolutionary attractor under the full mechanistic model, rather than a stochastic or contingent outcome.

